# Overlapping representation of primary tastes in a defined region of the gustatory cortex

**DOI:** 10.1101/114454

**Authors:** Max L. Fletcher, M. Cameron Ogg, Lianyi Lu, Robert J. Ogg, John D. Boughter

## Abstract

Two-photon imaging was used to examine taste responses in neurons in a region of gustatory cortex defined by thalamic input. This area contained an overlapping representation of primary tastes, with neurons that were either narrowly or broadly responsive to taste stimuli. Analysis demonstrates that activity in the neuronal population in this area yields information about both taste quality and hedonics.

## Introduction

Gustatory cortex (GC) plays an important role in the generation and maintenance of taste-related behaviors, including neophobia, taste aversion and other types of associative leaning (Braun et al., 1972; Lin et al., 2009; Samuelsen et al., 2012; Schier et al., 2014; Schier et al., 2016). Underlying these processes, cortical neurons encode information about both taste quality and hedonic valence (i.e. appetitive vs. aversive) (Yamamoto et al., 1985a, b; Katz et al., 2001; Jezzini et al., 2013). Yet there is still considerable debate as to how information about taste quality (i.e. sweet, salty, bitter and sour) and hedonic value are spatially represented and organized across GC. The primary gustatory cortical area is organized within the insular cortex, which in rodents is externally located on the ventral lateral brain surface, bisected by the middle cerebral artery (MCA) (Norgren and Wolf, 1975; Accolla et al., 2007; Chen et al., 2011; Kida et al., 2015).

Previous studies in anesthetized rats using physiological or intrinsic imaging techniques indicated that while basic taste stimuli tended to evoke distinctive spatial patterns across GC, there was substantial overlap, to the point that no one area was specific to a particular taste quality (Yamamoto et al., 1985b; Accolla et al., 2007). A similar finding was described in humans using functional magnetic resonance imaging (Schoenfeld et al., 2004). These findings contrast a recent 2-photon (2P) imaging study in mice (Chen et al., 2011), where large clusters of neurons responding only to a single taste quality, including a posterior bitter-responsive region and an anterior sweet-responsive region, were separated by sparsely-responsive regions across the cortical surface. Interestingly, there is evidence that spatial neuronal heterogeneity in present in some other sensory cortices. For example, in mouse piriform (olfactory) cortex, odorants of different classes evoke overlapping distributions (Stettler and Axel, 2009). In layer M/MI of mouse somatosensory cortex, whisker receptive fields are, to a degree, overlapping and scattered among columns (Clancy et al., 2015).

To gain further insight into the significance of spatial patterns of taste quality, we used 2P-imaging techniques to image taste responses to basic stimuli in an area of GC in mice immediately posterior to the MCA. This area was chosen due to the greatest probability of overlap among taste qualities, based on the rat studies mentioned above. We used a virally-expressed calcium reporter, GCaMP6s, shown to be sensitive enough to detect single action potentials from cortical cells *in vivo* with near 100% reliability (Chen et al., 2013). All recordings were performed from imaging windows located within an anatomically determined region of GC delineated by the MCA and bifurcation of the caudal rhinal vein. As the gustatory region of insular cortex can also be defined by taste thalamic input (Cechetto and Saper, 1987; Allen et al., 1991; Shi and Cassell, 1998; Nakashima et al., 2000) we also investigated anterograde labeling in this region following tracer injection into the gustatory subnucleus of the thalamus (VPMpc), including in some of the mice used for 2P imaging.

## Materials and Methods

### Animals and imaging surgery

Adult male and female C57BL/6J mice (Jackson Laboratories, Bar Harbor, ME) were used. All experimental protocols were approved by the University of Tennessee Institutional Animal Care and Use Committee. Mice were anesthetized with urethane (2mg/kg, ip). To reduce nasal secretions, mice also received i.p. injections of the blood brain barrier-impermeant muscarinic receptor antagonist, methyl scopolamine, at 0.05 mg/kg. Once mice were fully anesthetized, a tracheotomy was performed by inserting and securing a small piece of PE tubing into the trachea. Mice were then placed in a custom stereotaxic apparatus. The skin overlying the dorsal skull was removed and a head bar was attached using dental cement. The head of the animal was then tilted 90 degrees to allow for installation of the optical imaging window. To expose gustatory cortex, a small incision was made between the ear and eye. Portions of the masseter and temporalis muscles were cut away. The temporal portion of the zygomatic arch was removed to expose the lateral surface of the skull. The skull overlying the intersection of the MCA and rhinal veins was removed using a dental drill to create a small (approximately 2mm x 2mm) window. The surface of the brain was flushed with Ringer’s solution and covered with a 1% agarose solution and topped with a glass coverslip. The coverslip was fixed into place with dental cement to create a small well. Following collection of imaging data, permanent ink was applied to the brain surface, and in one mouse, an iontophoretic injection of 5% Fluorogold was made into the cortex at the recording site (these procedures were used to aid histological reconstruction). Mice were transcardially perfused with 4% paraformaldehyde, and brains were removed, post fixed, cryoprotected, and subsequently cut into coronal sections (40 μm) and examined for fluorescent labeling.

### Viral transfection and tracing

For viral transfections, mice were anesthetized with Ketamine/Xylazine (100 mg/kg and 10 mg/kg, ip) and placed into a stereotaxic head holder. For each animal, approximately 500 nL of AAV1.Syn.GCaMP6s (Penn Vector Core) was infused via picospritzer into the insular cortex (injection pipette was placed relative to bregma: anterior-posterior: +1.5mm, lateral: 3.7mm, depth: 2.1mm) through a craniotomy. Following infusion, mice were allowed to recover in their home cages for at least three weeks before imaging. For thalamic tracing experiments, 450 nL of tracer (AAV1.CB7.CI.mCherry; additional, non-imaging experiments used Microruby dextran, 3000 MW) was infused into the gustatory area of the thalamus (pipette was placed relative to bregma: anterior-posterior: −1.8mm, lateral: 0.6 mm, depth: 4.2 mm), which is located within the medial parvicellular portion of the ventral posterior medial division (VPMpc).

### Tastant delivery

Once mice were situated under the microscope, the oral cavity was gently held open via a loop of suture placed over the lower incisor. Whole mouth stimulation with water and stimuli representing four basic taste qualities (0.5 M sucrose, 0.3 M NaCl, 0.02 M citric acid, and 0.01 M QHCl, presented at room temperature) was delivered through a small length of PE tubing inserted into the mouth and connected to a manifold, which allowed for switching between water and solutions. During trials, mice received a constant flow (rate = 0.25 ml/s) consisting of an 8 s water presentation, a 10 s tastant presentation and a 12 s water rinse. After the trial, the oral cavity was rinsed with water for an additional 20 s and then allowed to recover for 1.5 minutes prior to the next trial. Stimuli were presented in random order.

### Optical imaging and analysis

Imaging was performed on a Zeiss 7MP 2-photon microscope equipped with a Zeiss 20x objective. Fluorescence images were collected at 2-4Hz at 512 x 512 pixel resolution. Imaging regions ranged from 200μm^2^ to 500μm^2^. Individual cells could easily be identified in resting fluorescence images. ROIs were manually drawn around all cells within an imaging region and raw fluorescence traces for the entire 40 seconds of each trial were collected off-line using Image J and analyzed using the R software package.

Analysis was performed in R Studio (Version 0.98.1103). All raw fluorescence traces were compiled and interpolated to the fastest frame rate (t=0.309 s) using the *spline* function. Traces were smoothed using a 3 frame rolling mean function. To quantify taste-specific responses, the taste-evoked change in fluorescence ΔF) from each trace was calculated by subtracting the 8s frame average during the pre-taste water presentation from a 5-frame average centered on the peak of the response generated during the taste presentation. The relative change in fluorescence (ΔF/F) was then calculated by dividing the taste-evoked change in fluorescence by the mean fluorescence. Responsive cells were defined as having a ΔF/F greater than that of the mean ± 2.5 SD of the pre-taste water presentation for each trial.

Entropy, a measure of breadth of tuning, was calculated for each neuron (Smith et al., 1979)

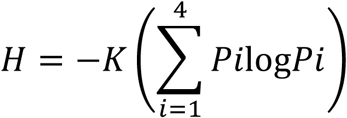

where *Pi* represents the response to each of the four taste stimuli and K is a scaling constant (1.661 for four stimuli). Values of entropy (*H*) close to zero indicate reponse to only a single stimulus (narrow tuning) while values close to 1.0 indicate response to all four stimuli (broad tuning). We also calculated an additional measure of tuning breadth: the noise-to-signal ratio (N/S) (Spector and Travers, 2005). This ratio is derived by dividing the response to the second best stimulus (the maximum noise elicited by sideband stimuli) by the response to the best stimulus (signal). Similar to *H,* this measure also ranges from 0.0 to 1.0.

For cluster analysis, all taste responses for each cell were normalized to the maximum taste-evoked response for that cell. Hierarchical cluster analysis and principle components analysis were then performed using JMP software. Mean normalized distances between cells of each cluster were calculated by dividing the distance between each cell of a cluster by the mean the distance between all the cells in the imaging window.

### Immunocytochemistry

Intraoral (IO) cannulas were affixed to the skull in some anesthetized mice following thalamic tracer (Microruby) injection. Polyethylene tubing was inserted through the right buccal mucosa, led along the lateral surface of the skull, and secured to the skull using dental acrylic. Three days after surgery mice underwent an adaptation procedure, receiving distilled water through the IO cannula (0.1 ml/min for 15 min) using a syringe pump in a round Plexiglas test chamber. After 3 consecutive days of this adaptation procedure, mice were intraorally infused with 1.5 ml of 0.003 M QHCl using the same methods and rate as the adaptation procedure. Two hours after the onset of IO stimulation, mice were anesthetized with Ketamine/Xylazine (100 mg/kg and 10 mg/kg, ip) and transcardially perfused with 4% paraformaldehyde. The brains were removed, post fixed, and cryoprotected; coronal sections of the brain were cut serially using a freezing microtome. Antigen expression, including c-Fos, was assessed using standard IHC procedures, with a rabbit polyclonal anti-c-Fos antibody (sc-52, Santa Cruz Biotechnology); M2-type muscarinic acetylcholine receptors were labeled using a rat monoclonal anti-M2 antibody (MAB 367, EMD Millipore). Primary antibody labeling was visualized with either fluorescent or non-fluorescent secondary antibodies (see our previous papers for detailed methods, (Savchenko and Boughter, 2011; Tokita and Boughter, 2016). Microscope sections were imaged using either a Leica (DMRXA2, Leica Microsystems) microscope equipped with a digital camera and imaging software, or with a confocal microscope (Zeiss 710).

### Brief-access taste behavior

Water-restricted C57BL/6J mice (n=7) were tested in a Davis MS-160 contact lickometer with a panel of taste stimuli (0.5 M sucrose, 0.3 M NaCl, 0.02 M citric acid, 0.01 M QHCl) and water. This method involved 2 days of training with water only, and 2 days of testing; procedures are based on previous studies by the authors (StJohn and Boughter, 2009; Saites et al., 2015). On the two test days, mice received 18 5-s trials, divided into 3 blocks consisting of a single presentation of each tastant and 2 presentations of water. Stimuli were ordered randomly within each block. Data were averaged across the two days of testing for each animal and are presented in the form of mean lick ratios (average licks to stimulus/average licks to water) for each stimulus.

## Results

We used 2P imaging of the virally-expressed calcium reporter GCaMP6s to study taste quality representation within GC neurons in cortical layers II/III, testing whether they possess specific responses and examining whether they tend to group together in space across the GC surface. We first verified the extent of GC in mice by making injections of the anterograde tracer microruby into the medial parvocellular portion of the ventral posterior medial division of the thalamus (VPMpc) (Figure 1C). In all mice (n=7), VPMpc injections consistently labeled thalamic fibers that ramified throughout all cortical layers of granular and disgranular cortex ranging from approximately −0.5 mm anterior to bregma to 1.5 mm posterior to bregma (Figure 1E). We also found consistent taste-evoked (QHCl) c-fos labeling in regions containing labeled thalamic fibers, further demonstrating the labeled area is GC (Figure 1G).

**Figure 1.**
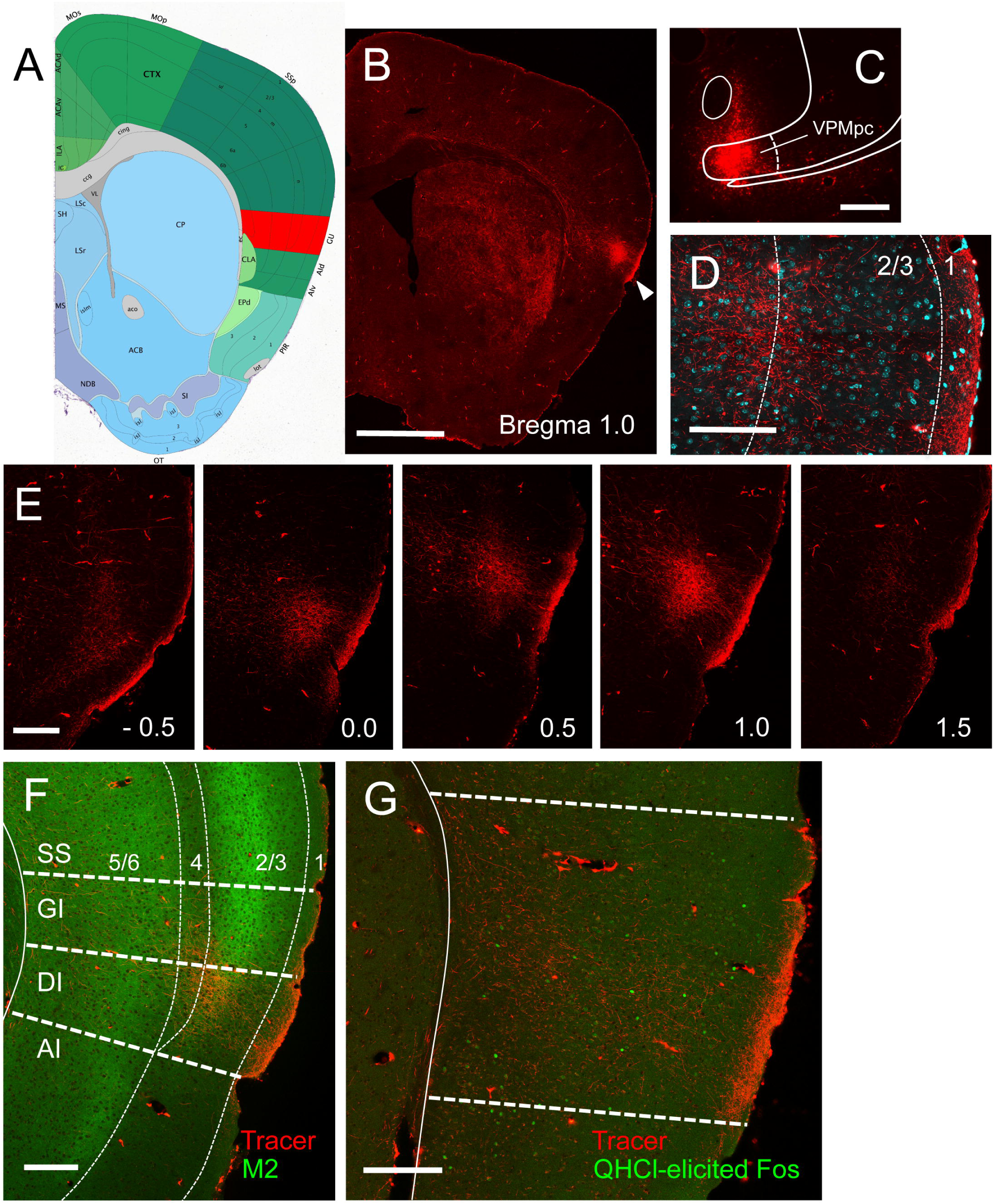
Delineation of GC with neural tracing and immunocytochemistry. Following taste-evoked imaging, mice were perfused transcardially, and in two mice the posterior edge of the MCA was marked directly on the brain surface using a blue dye. Serial sectioning of these brains (not shown) indicated that the MCA was located approximately at 1.0 mm anterior to bregma, a level corresponding to images 44-45 of the P56 Coronal Reference Atlas, part of the Allen Brain Atlas (ABA; http://www.brain-map.org/). Image 45 is shown in (**A**), with GC (labeled GU on the image) as identified by ABA colored in red. Several mice (n = 7) were used in neural tracing studies to further explore and corroborate the boundaries of GC. An anterograde tracer (microruby) was injected unilaterally into the gustatory thalamus (VPMpc) of B6 mice; several weeks later terminal labeling was examined in GC. **B**. Whole-brain section shows terminal labeling concentrated in the ipsilateral cerebral hemisphere in a dorsal-lateral position (arrowhead), and at an anterior-posterior level (approximately 1.0 mm anterior to bregma), consistent with the designation of GC according to ABA. **C**. Tracer Injection site in VPMpc. **D**. Magnified view of terminal labeling (red) interposed with cellular stain (DAPI; cyan) shows diffuse thalamic input, with concentrations in cortical layers 1,3 and 4-5. **E**. Sequential sections from the same mouse show labeling from caudal – rostral levels. Terminals were found at each of these levels, shown in 0.5 mm increments from −0.5 to 1.5 mm anterior to bregma, but labeling was most robust from about 0.0 to about 1.2 mm. This corresponds to the center of the GC according to ABA. **F**. Staining of cortex with an antibody to the muscarinic M1-type receptor (green) is noteworthy in that it is concentrated in layers 3 and 5 of 6-layer neocortex; the expression in layer 2 dissipates ventrally as the granular insular cortex gives way to the dysgranular insular cortex, which is cytoarchitecturally characterized by the gradual disappearance of layer 4 (the granular layer). Contrasting M2 expression in GC with thalamic tracing shows that the thalamic input (red) is largely located in the dysgranular insular cortex, a finding that is consistent with previous studies in rats (e.g.Allen et al., 1991). **G**. Taste-evoked Fos expression (0.003 M QHCl intraorally delivered to an awake mouse) also overlaps with the thalamic terminal field in GC. Scale bars: B = 1 mm, C = 0.5 mm, D = 0.1 mm, E-G = 0.2 mm. Pictures B, C, and E are from the same mouse; pictures D and F are from another, and picture G is from another. All boundaries superimposed on images are approximate.

As previous imaging studies focused on an anatomically defined region of GC delineated by the MCA and bifurcation of the caudal rhinal veins (Accolla et al., 2007; Chen et al., 2011), we targeted our viral injections to this region (corresponding to +1.5mm from bregma) (Figure 2A, B, D). 2P imaging in these mice reveal large populations of labeled cells within layer II/III of our imaging window. In a subset of these mice, we also injected AAV1.CB7.CI.mCherry to label thalamic fibers. As can be seen in Figure 2C, we found dense overlap between thalamic fibers (red) and GCaMP-labeled GC cells (green) further verifying imaging region is located within GC.

**Figure 2.**
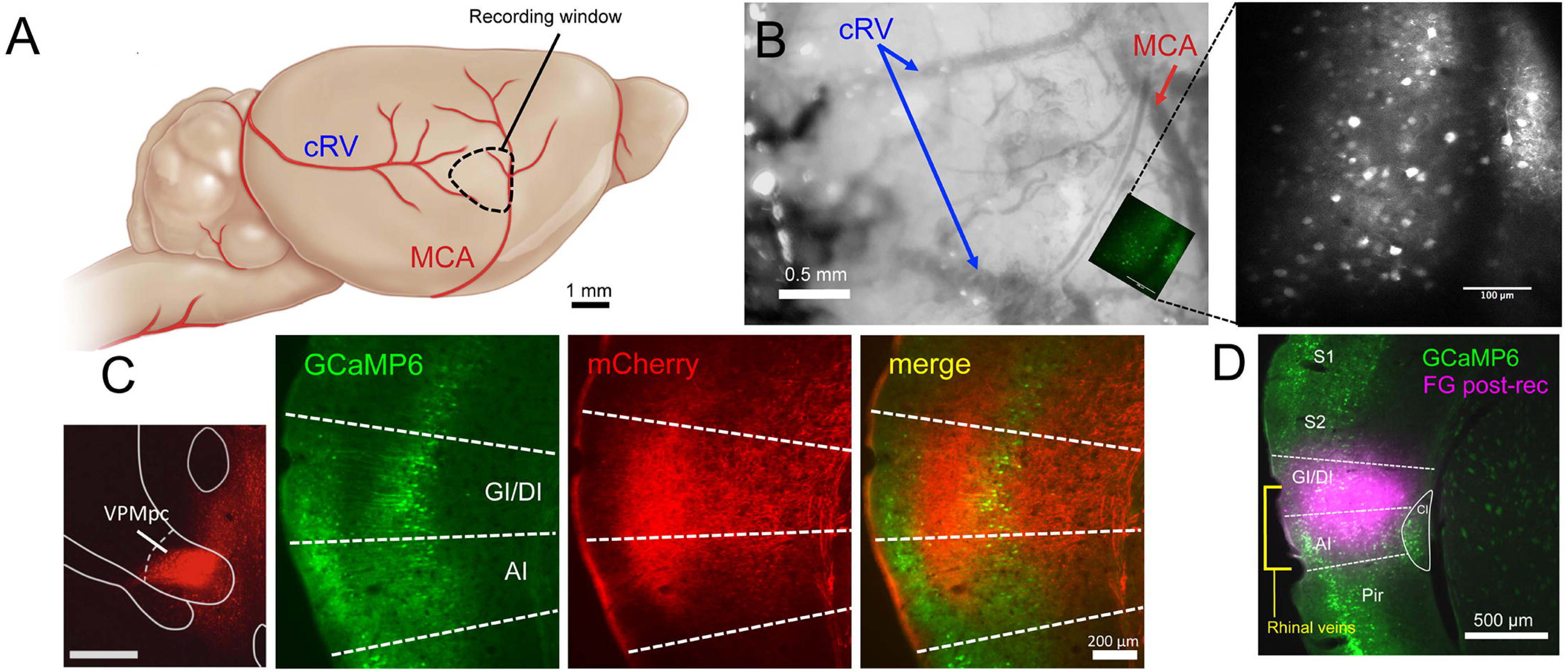
Population recording from an anatomically defined region of GC. **A**. Diagram depicting region of recordings based on vasculature. **B**. Example image taken from a recording window with one imaging field superimposed. Right: 2-photon image of example field showing labeled neurons. **C**. The GCaMP-visualized neurons were verified to be within GC by the presence of anterograde mCherry labeled VPMpc thalamic fibers. **D**. In some animals, following imaging, FluoroGold was injected into the recording area to further verify recordings were from GC neurons.

In seven mice, we recorded significant taste-evoked activity in 88% of examined neurons (783 of 891 cells in 24 microscope fields). For each field, responses were recorded from all visible cells allowing us to generate a taste profile and spatial location for each cell. Individual cell taste responses were repeatable across taste presentations (Figure 3A) and the overall percentage of responding cells did not change as stricter response criteria were applied (see methods) (Figure 3B). Both example traces and population data show a large number of cells responding to each of the tastes, as well individual cells responding to multiple tastes (Figure 3C-E). Significant taste responses in individual cells ranged from 9.8-492.0% ΔF/F (mean response: 32.9±0.8% ΔF/F). The percentage of cells with best responses to each tastant was similar (C: 29%, N: 19%, Q: 25%, S: 26%) (Figure 3F). The percentage of cells responding to each tastant (C: 60%, N: 54%, Q: 51%, S: 49%)(Figure 3G) and the overall mean responses to each tastant (C: 30.3±1.3%, N: 33.7±1.7%, Q: 28.6±1.4%, S: 39.0±2.2%) (Figure 3H) were similar, with substantial overlap in this region.

**Figure 3.**
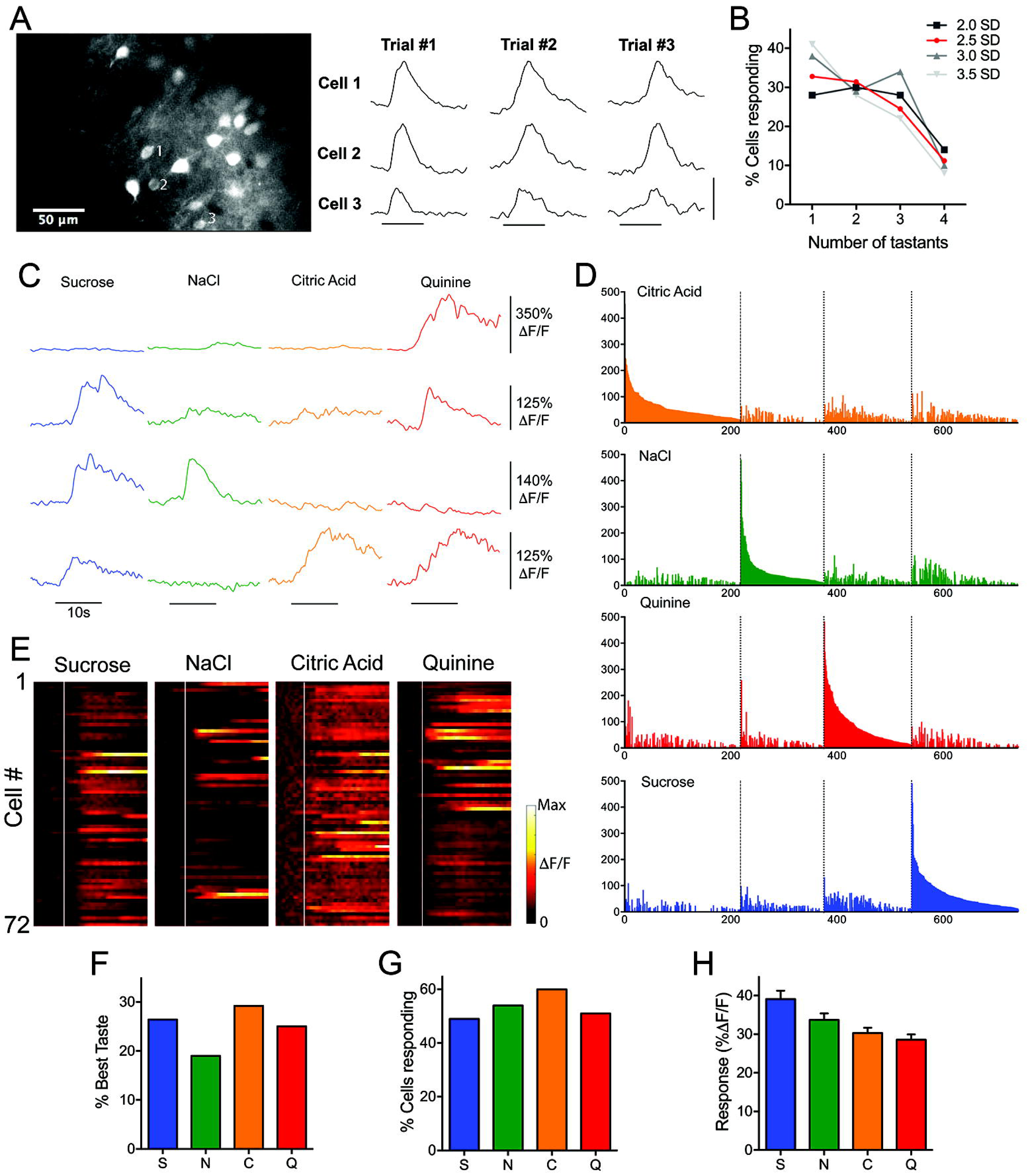
**A**. Resting fluorescence imaging of layer M/MI GC neurons. Fluorescence traces show multiple taste presentations, separated by several minutes, evoke similar responses from the three labeled neurons. Horizontal scale bar = 5 sec, Vertical scale bar = 200% ΔF/F. **B**. Individual cell response thresholds were set at 2.0, 2.5, 3.0, and 3.5 standard deviations (SD) and the percentage of cells responding to only one, two, three, or four tastants were compared. All other thresholds (grays) produced highly similar percentages of cells as the 2.5SD (red) used study. **C.** Example traces from four different GC neurons to taste application (black bar). **D**. Normalized taste-evoked responses from all cells taken from the field shown in Figure 2B. White line depicts onset of taste delivery. Individual neuron responses vary in their selectivity and temporal response. **E**. All responsive neurons grouped into best-stimulus categories (colors) and arranged in descending order of response magnitude to that stimulus. **F.** Percentage of cells by preferred taste. **G.** Similar percentages of cells significantly responding to each taste. **H.** Mean fluorescence change evoked by each taste across the cell population.

To investigate population coding, we pooled all responsive cells from all mice and performed multivariate analysis of normalized taste responses. Hierarchical cluster analysis of the responses identified nine distinct neural taste profiles (Figure 4A, B, Table 1. Importantly, cells representing each cluster could be found in all mice. For each cluster, we calculated the mean entropy (Figure 4C, Table 1) (Smith et al., 1979) and mean noise-to-signal (N/S) value (Figure 4D, Table 1) (Spector and Travers, 2005). Based on this, the cell clusters comprised of two taste quality groups, with four singly-tuned clusters representing each individual tastant and five broadly-tuned clusters preferring different combinations of tastants. Using this classification, we found a majority (65%) of cells to be narrowly tuned rather than broadly tuned (Figure 4E).

**Figure 4.**
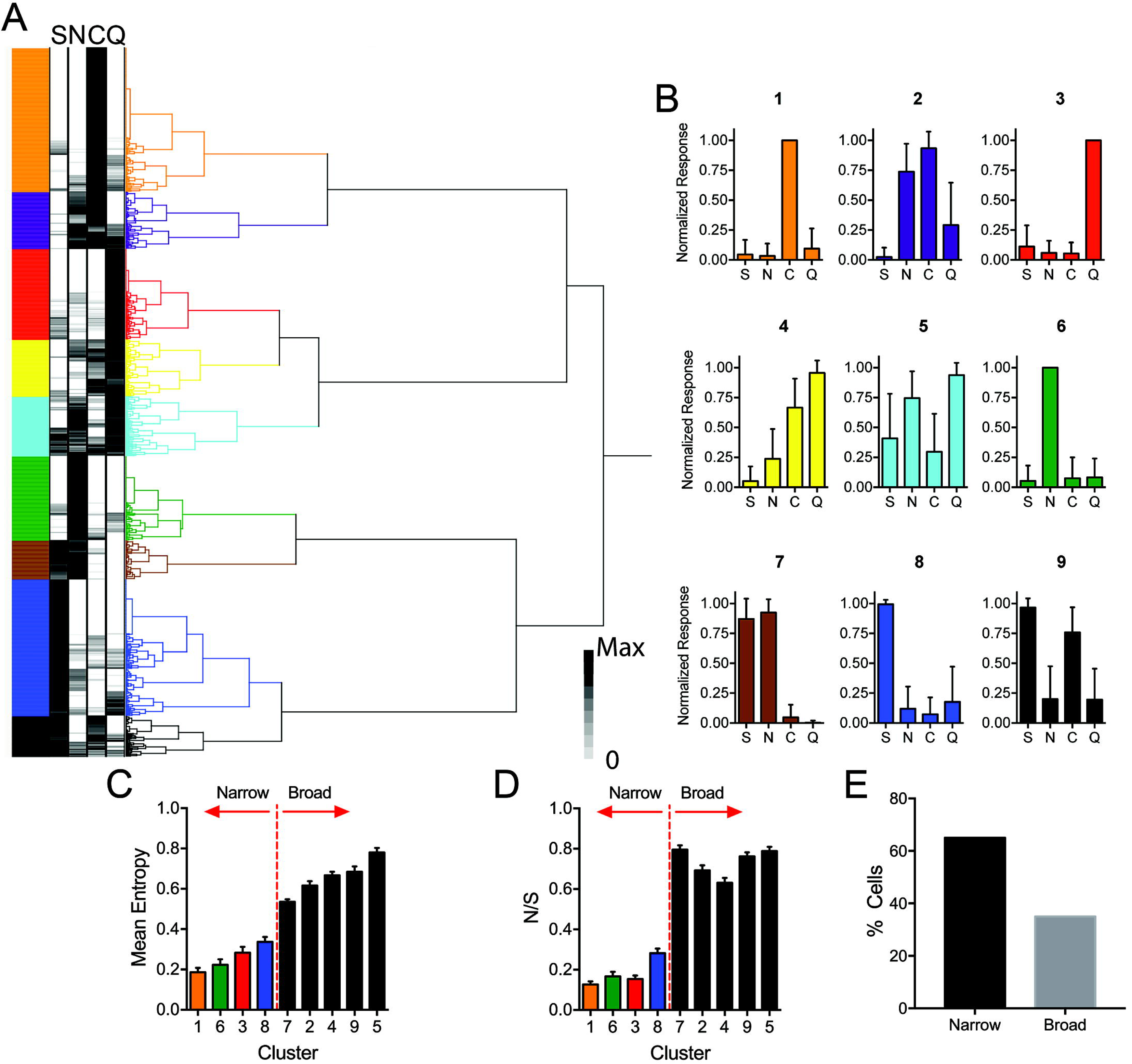
**A**. Hierarchical cluster analysis reveals groups of cells with distinct taste response profiles. **B**. Normalized taste response profiles (mean±SD) for each cluster in A reveal both singly tuned and more broadly tuned cell types. Entropy (**C**) and Noise-to-Signal (**D**) analysis demonstrate narrowly and broadly tuned cell clusters. **E**. Percentage of strongest (top 10% by max response) responding cells according to cluster.

Principal components analysis demonstrated robust separation of primary taste qualities along the first three components axes (Figure 5A). Further, hedonic character separated along the first component effectively separating aversive stimuli (citric acid and quinine) from neutral (NaCl) or appetitive tastes (sucrose). Behavioral taste preference data using the same taste concentrations displayed similar patterns, with mean lick ratios relative to water differing significantly according to each taste quality (S: 1.34±0.10, N: 0.99±0.17, C: 0.46±0.07, Q: 0.20±0.03) (One-way ANOVA, F _[3,24]_ = 23.2, p < 0.0001) (Figure 5B). One sample t-tests revealed water-restricted mice preferred sucrose (t(6)=3.14, p=0.02) and avoided citric acid (t(6)=7.82, p<0.001) and quinine (t(6)=23.56, p<0.001) while displaying no preference for NaCl (t(6)=0.04, p=0.97).

**Figure 5.**
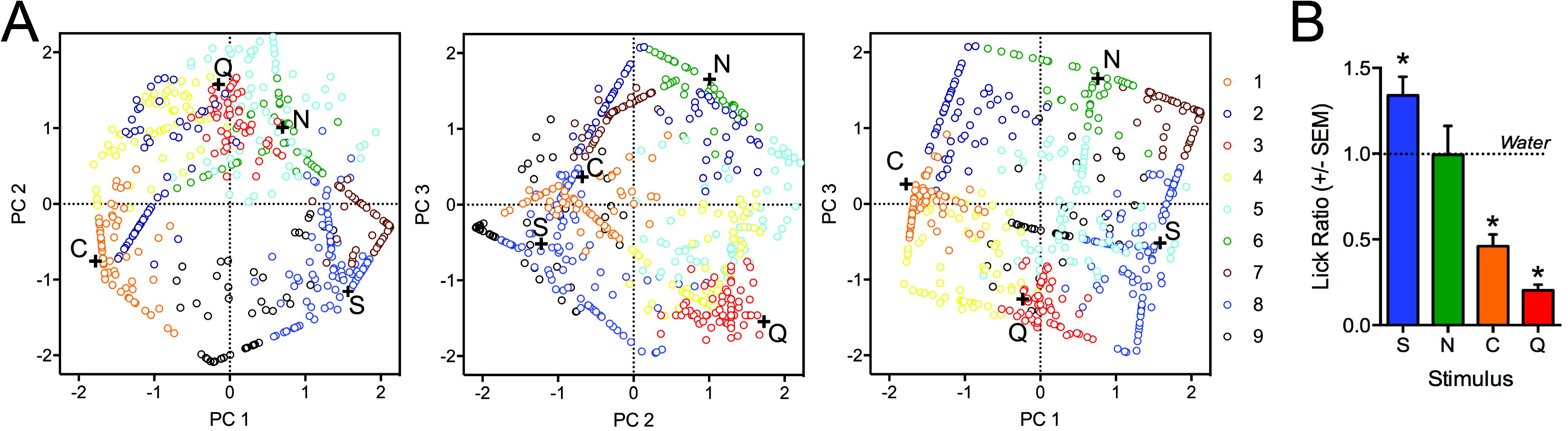
**A**. Two-dimensional plots of the first three principal components (accounting in for 91% of the total variance) relative to one another reveal that the population data effectively and separately encodes each tastant in coding space. When plotted along the first principal component, the taste clearly separate along a hedonic spectrum from appetitive to aversive. In all graphs, the individual cell positions are plotted and color coding according to their cluster to demonstrate how each cluster contributes to the coding of each taste. **B.** Mean lick ratios relative to water differ according to each taste quality. Overall, water-restricted mice preferred sucrose and avoided citric acid and quinine while displaying no preference for NaCl compared to water. Asterisks above bars indicate significantly different groups. Error bars represent SEM.

Finally, to investigate the spatial distribution of cells along the surface of GC based on quality response and tuning, we determined the location of each cell within each imaging field. Response maps were constructed on a Cartesian grid, with each cell marked by response cluster (Figure 6A) or tuning breadth (broad or narrow) (Figure 6B). Across all imaging windows, no obvious grouping of cells was seen based on cluster number. Further, for each cluster, the normalized pair-wise distances between cells was not significantly different than one (Wilcoxon signed rank test, theoretical median of 1; p<0.05 for all clusters), suggesting that cells belonging to individual clusters do not tend to be spatially grouped within a given field (mean values: cluster 1=0.99±0.09; cluster 2=0.76±0.10; cluster 3=0.92±0.05; cluster 4=0.95±0.10; cluster 5=0.97±0.13; cluster 6=1.01±0.09; cluster 7=0.85±0.09; cluster 8=1.07±0.07; cluster 9=0.79±0.10) (Figure 6C). However, when cells were grouped according to entropy, broadly tuned cells were found to be closer together on average than narrowly tuned cells (mean values: narrow=1.01±0.04; broad=0.87±0.05) (Mann Whitney test, p=0.009) and significantly less than the mean distance between all cells (Wilcoxon signed rank test, theoretical median of 1; p=0.01) (Figure 6D), suggesting some clustering of broadly tuned cells within GC.

**Figure 6.**
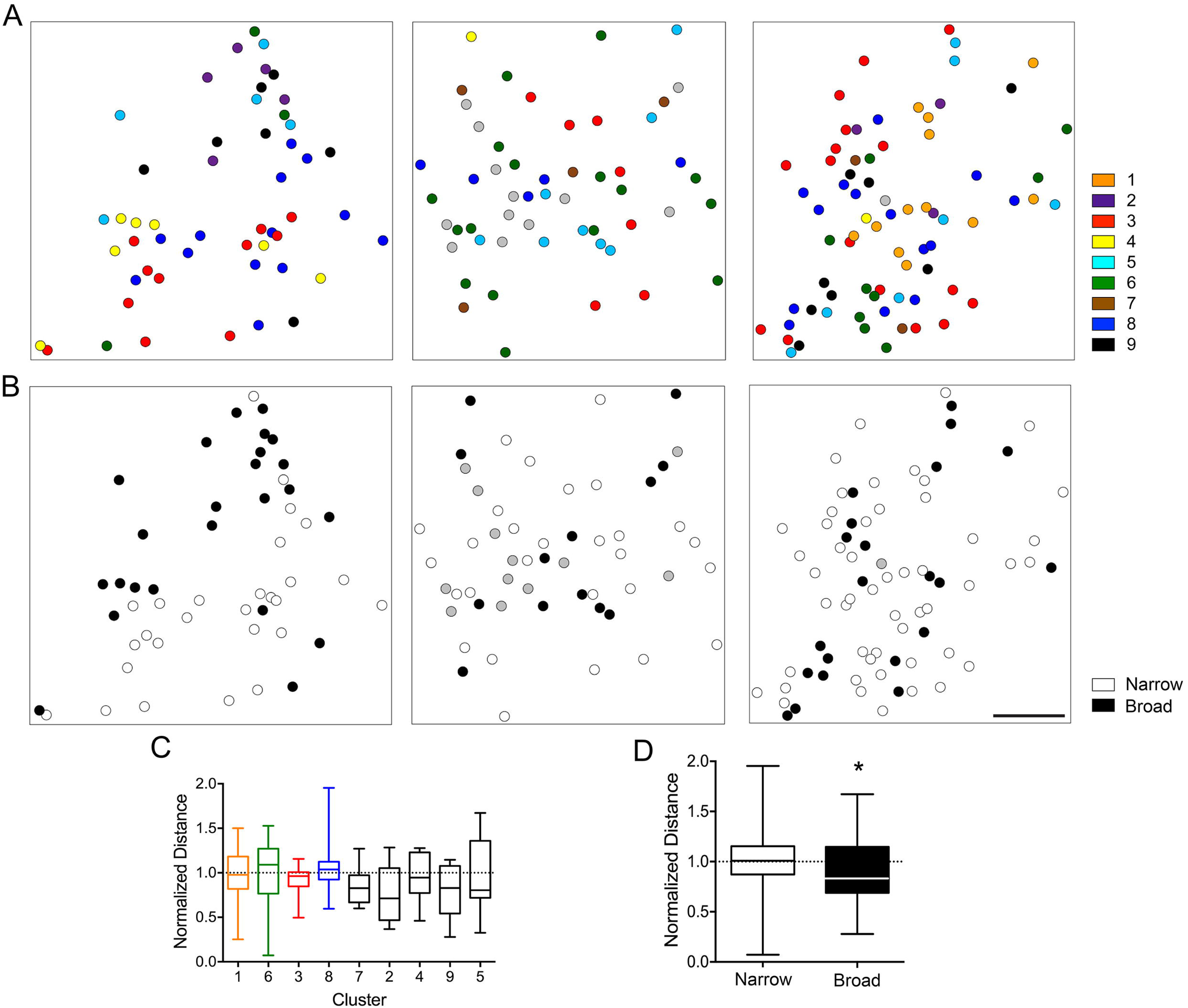
Examples of the spatial distribution of neurons within individual fields of GC in different mice based on cluster (**A**) and tuning breadth (**B**). **C**. Percent of total cells belonging to each cluster. **D**. Normalized distance between cells of each cluster. **E**. Normalized distance between narrowly tuned and broadly tuned cells. Bars represent minimum and maximum distances.

## Discussion

### Coding of taste quality and hedonics in cortical neurons

Cortical taste neurons varied in their responses to stimuli representing the four basic tastes. Neurons were found that responded to just one of the four primary tastes, while other neurons were more broadly tuned, responding to two, three or even all four stimuli. Nearly all gustatory physiology studies published over the last half century have reported the existence of both “specialist” and “generalist” taste-responsive cells, from peripheral sensory neurons to the cortex, in studies with either anesthetized or awake and behaving animals (Spector and Travers, 2005; Simon et al., 2006; Carleton et al., 2010). Using hierarchical cluster analysis, we identified roughly comparable numbers of narrowly tuned taste neurons, or specialists, for each of the basic taste stimuli. Collectively, specialists accounted for 64% of all taste-responsive cells. Interestingly, this ratio resembles that found for geniculate ganglion taste neurons in two recent imaging papers (Barretto et al., 2015; Wu et al., 2015). Overall neuronal selectivity can also be inferred by breadth-of-tuning measures such as entropy; in these experiments overall mean entropy was 0.40 ± 0.01 (SEM). Studies of taste brainstem physiology in C57BL/6J mice reveal higher entropies (i.e. more broad tuning), 0.73 and 0.64 for the nucleus of the solitary tract and parabrachial nucleus, respectively (Lemon and Margolskee, 2009; Tokita and Boughter, 2016). There does not appear to be greater convergence of taste quality information into individual cells in the cortex vs. lower areas, although evidence for sharpening of taste profiles in the former region in rat studies at least is equivocal (Spector and Travers, 2005).

What then, are the functional implications of specialist and generalist taste cell types in mouse GC? There is some evidence, albeit limited, that variation in breadth-of-tuning could reflect different cell types, i.e. pyramidal cells vs. interneurons (Yokota et al., 2011). Tuning profiles may also correlate with cortical depth or projection. Projection cells, located in GC in rats primarily in layer V/VI, target other parts of cortex, including ipsilateral GC and orbitofrontal cortex, as well as subcortical structures such as thalamus, amygdala, and brainstem (Maffei et al., 2012). Our imaging responses were collected within layer II/III. There was actually evidence of spatial clustering of broadly tuned neurons (but not taste quality *per se*) in our data, possibly indicating a shared function of such cells. Interestingly, our data adhered well to the specific-to-general cell-assembly model recently described by Xie et al. (2016) for other cortical areas. In this model, distinct stimuli activate pyramidal cells according to power-of-two permutation based logic, isuch that neural networks are organized to respond to relevant and possibly all combinations of stimuli, including specialists and generalists. Specific cells are predicted to be overrepresented in superficial cortical layers, as was found in our study.

The diverse connection pattern apparent in combinations of neuronal responder types offers increased computational power and flexibility (Xie et al., 2016), which are ideal features for GC, where stimulus processing is linked to attentional states and taste-based leaning (Fontanini and Katz, 2008; Samuelsen et al., 2012; Schier et al., 2016). Awake multi-electrode recordings in rats indicate that information reflecting multiple facets of taste, including quality and hedonic valence, are encoded in the activity of ensembles of cortical cells (Katz et al., 2001; Jezzini et al., 2013). Although we are not analyzing temporality of spiking, making direct comparison to these studies difficult, principle components analysis of our data indicated that both quality and hedonic character are encoded in the activation patterns of all cell types. For hedonics, this reflects the degree of convergence of aversive, appetitive or neutral stimuli in broadly tuned cells.

### Topography of taste quality in GC

The issue of whether there is a spatial map of taste quality within particular brain areas has received a fair amount of study, even in human taste GC (Singh et al., 2011). GC comprises a subregion of so-called insular cortex, and in rodents is located on the lateral ventral surface of the brain. Its spatial and functional characteristics have been much better defined in rats than mice. According to the Allen Brain Atlas (ABA), mouse GC extends for ~ 2.4 mm along the anterior-posterior (AP) axis, corresponding to about +2.0 to −0.2 from bregma (Paxinos and Franklin, 2001). We imaged taste cell responses just caudal to the MCA, which we estimate lies on the external brain surface at about +1.0 mm from bregma (although in rats the location of MCA with respect to actual brain landmarks has been shown to be variable; (Hashimoto and Spector, 2014; Kida et al., 2015). Abundant input from the taste thalamus (VPMpc) was found in this region, in a dorsal-ventral location similar to the designation of GC in ABA, and corresponding primarily to the dysgranular insular cortex (characterized by the attenuation of layer 4).

Our imaging responses provide compelling evidence in mice that the rostral-caudal “center” of GC contains overlapping representations of taste quality, challenging notions of labeled-line coding via segregated, quality-specific cell clusters (Chen et al., 2011). This overlap among basic taste qualities in the center region is similar to results gained using other approaches in anesthetized rats, including wide-field optical imaging (Accolla et al., 2007; Accolla and Carleton, 2008) as well as single unit *in vivo* recordings (Yamamoto et al., 1985b). Moreover, this area is targeted in most awake recording studies, linking taste responses to behavioral function (Katz et al., 2001; Fontanini and Katz, 2008; Samuelsen et al., 2012; Maier and Katz, 2013). Lesions centered in this area have effects on quinine and salt, but not sucrose, sensitivity and discrimination (Bales et al., 2015; Blonde et al., 2015).

The tension between our results and the Chen et al. study (2011) may be due to several different factors. The methodology is somewhat different, as the current study utilized virally-expressed GCaMP6s rather than a bulk-loaded AM dye (used in the previous study), allowing for greater penetration into cortical layers II/III. It should also be noted that the Chen et al. study investigated areas more anterior or posterior to the region we focused on; it is possible that greater concentrations of particular best-response cell types are found at the rostral or caudal extremes of gustatory cortex, while there is more mixing in the center, as in the rat studies (Yamamoto et al., 1985b; Accolla et al., 2007). It must be emphasized, however, that in these previous rat mapping studies, any such biasing at the extremes corresponded to a gradation of numbers of particular best-taste cell types, rather than discrete and absolute spatial clustering (Yamamoto et al., 1985b; Accolla et al., 2007). Indeed, gradients of especially bitter and sweet-best neurons have been described in other central gustatory areas such as the nucleus of the solitary tract (NTS) or parabrachial nucleus (PBN), where they most likely reflect the differential expression of taste receptors (Gilbertson and Boughter, 2003) in the oral cavity by cranial nerves (Travers and Norgren, 1995; Karimnamazi et al., 2002; Geran and Travers, 2006; Yokota et al., 2011; Tokita and Boughter, 2016).

## Author Contributions

M.L.F. and J.D.B designed the research project; M.L.F., J.D.B., L.L. performed the experiments. M.L.F., M.C.O., and R.J.O analyzed the data. M.L.F. and J.D.B wrote the manuscript.

## Acknowledgements

The authors gratefully acknowledge the assistance of Stephanie Staszko and Joseph Callaway with aspects of this research. This study was supported by NIH grant NIDCD 015202 to M.L.F. and J.D.B. and the Pew Biomedical Scholars Program Grant to M.L.F.

## References

Accolla R, Carleton A (2008) Internal body state influences topographical plasticity of sensory representations in the rat gustatory cortex. Proc Natl Acad Sci U S A 105:4010-4015.

Accolla R, Bathellier B, Petersen CC, Carleton A (2007) Differential spatial representation of taste modalities in the rat gustatory cortex. J Neurosci 27:1396-1404.

Allen GV, Saper CB, Hurley KM, Cechetto DF (1991) Organization of visceral and limbic connections in the insular cortex of the rat. J Comp Neurol 311:1-16.

Bales MB, Schier LA, Blonde GD, Spector AC (2015) Extensive Gustatory Cortex Lesions Significantly Impair Taste Sensitivity to KCl and Quinine but Not to Sucrose in Rats. PLoS One 10:e0143419.

Barretto RP, Gillis-Smith S, Chandrashekar J, Yarmolinsky DA, Schnitzer MJ, Ryba NJ, Zuker CS (2015) The neural representation of taste quality at the periphery. Nature 517:373-376.

Blonde GD, Bales MB, Spector AC (2015) Extensive lesions in rat insular cortex significantly disrupt taste sensitivity to NaCl and KCl and slow salt discrimination learning. PLoS One 10:e0117515.

Braun JJ, Slick TB, Lorden JF (1972) Involvement of gustatory neocortex in the learning of taste aversions. Physiol Behav 9:637-641.

Carleton A, Accolla R, Simon SA (2010) Coding in the mammalian gustatory system. Trends Neurosci 33:326-334.

Cechetto DF, Saper CB (1987) Evidence for a viscerotopic sensory representation in the cortex and thalamus in the rat. J Comp Neurol 262:27-45.

Chen TW, Wardill TJ, Sun Y, Pulver SR, Renninger SL, Baohan A, Schreiter ER, Kerr RA, Orger MB, Jayaraman V, Looger LL, Svoboda K, Kim DS (2013) Ultrasensitive fluorescent proteins for imaging neuronal activity. Nature 499:295-300.

Chen X, Gabitto M, Peng Y, Ryba NJ, Zuker CS (2011) A gustotopic map of taste qualities in the mammalian brain. Science 333:1262-1266.

Clancy KB, Schnepel P, Rao AT, Feldman DE (2015) Structure of a single whisker representation in layer 2 of mouse somatosensory cortex. J Neurosci 35:3946-3958.

Fontanini A, Katz DB (2008) Behavioral states, network states, and sensory response variability. J Neurophysiol 100:1160-1168.

Geran LC, Travers SP (2006) Single neurons in the nucleus of the solitary tract respond selectively to bitter taste stimuli. J Neurophysiol 96:2513-2527.

Gilbertson TA, Boughter JD Jr., (2003) Taste transduction: appetizing times in gustation. Neuroreport 14:905-911.

Hashimoto K, Spector AC (2014) Extensive lesions in the gustatory cortex in the rat do not disrupt the retention of a presurgically conditioned taste aversion and do not impair unconditioned concentration-dependent licking of sucrose and quinine. Chem Senses 39:57-71.

Jezzini A, Mazzucato L, La Camera G, Fontanini A (2013) Processing of hedonic and chemosensory features of taste in medial prefrontal and insular networks. J Neurosci 33:18966-18978.

Karimnamazi H, Travers SP, Travers JB (2002) Oral and gastric input to the parabrachial nucleus of the rat. Brain research 957:193-206.

Katz DB, Simon SA, Nicolelis MA (2001) Dynamic and multimodal responses of gustatory cortical neurons in awake rats. J Neurosci 21:4478-4489.

Kida I, Enmi J, Iida H, Yoshioka Y (2015) Asymmetrical intersection between the middle cerebral artery and rhinal vein suggests asymmetrical gustatory cortex location in rodent hemispheres. Neurosci Lett 589:150-152.

Lemon CH, Margolskee RF (2009) Contribution of the T1r3 taste receptor to the response properties of central gustatory neurons. J Neurophysiol 101:2459-2471.

Lin JY, Roman C, St Andre J, Reilly S (2009) Taste, olfactory and trigeminal neophobia in rats with forebrain lesions. Brain research 1251:195-203.

Maffei A, Haley M, Fontanini A (2012) Neural processing of gustatory information in insular circuits. Curr Opin Neurobiol 22:709-716.

Maier JX, Katz DB (2013) Neural dynamics in response to binary taste mixtures. J Neurophysiol 109:2108-2117.

Nakashima M, Uemura M, Yasui K, Ozaki HS, Tabata S, Taen A (2000) An anterograde and retrograde tract-tracing study on the projections from the thalamic gustatory area in the rat: distribution of neurons projecting to the insular cortex and amygdaloid complex. Neurosci Res 36:297-309.

Norgren R, Wolf G (1975) Projections of thalamic gustatory and lingual areas in the rat. Brain research 92:123-129.

Saites LN, Goldsmith Z, Densky J, Guedes VA, Boughter JD, Jr., (2015) Mice perceive synergistic umami mixtures as tasting sweet. Chem Senses 40:295-303.

Samuelsen CL, Gardner MP, Fontanini A (2012) Effects of cue-triggered expectation on cortical processing of taste. Neuron 74:410-422.

Savchenko VL, Boughter JD Jr.,(2011) Regulation of neuronal activation by Alpha2A adrenergic receptor agonist. Neurotox Res 20:226-239.

Schier LA, Blonde GD, Spector AC (2016) Bilateral lesions in a specific subregion of posterior insular cortex impair conditioned taste aversion expression in rats. J Comp Neurol 524:54-73.

Schier LA, Hashimoto K, Bales MB, Blonde GD, Spector AC (2014) High-resolution lesion-mapping strategy links a hot spot in rat insular cortex with impaired expression of taste aversion learning. Proc Natl Acad Sci U S A 111:1162-1167.

Schoenfeld MA, Neuer G, Tempelmann C, Schussler K, Noesselt T, Hopf JM, Heinze HJ (2004) Functional magnetic resonance tomography correlates of taste perception in the human primary taste cortex. Neuroscience 127:347-353.

Shi CJ, Cassell MD (1998) Cortical, thalamic, and amygdaloid connections of the anterior and posterior insular cortices. J Comp Neurol 399:440-468.

Simon SA, de Araujo IE, Gutierrez R, Nicolelis MA (2006) The neural mechanisms of gustation: a distributed processing code. Nat Rev Neurosci 7:890-901.

Singh PB, Iannilli E, Hummel T (2011) Segregation of gustatory cortex in response to salt and umami taste studied through event-related potentials. Neuroreport 22:299-303.

Smith DV, Travers JB, Van Buskirk RL (1979) Brainstem correlates of gustatory similarity in the hamster. Brain Res Bull 4:359-372.

Spector AC, Travers SP (2005) The representation of taste quality in the mammalian nervous system. Behav Cogn Neurosci Rev 4:143-191.

St John SJ, Boughter JD, Jr.,(2009) Orosensory responsiveness to and preference for hydroxide-containing salts in mice. Chem Senses 34:487-498.

Stettler DD, Axel R (2009) Representations of odor in the piriform cortex. Neuron 63:854-864.

Tokita K, Boughter JD, Jr., (2016) Topographic organizations of taste-responsive neurons in the parabrachial nucleus of C57BL/6J mice: An electrophysiological mapping study. Neuroscience 316:151-166.

Travers SP, Norgren R (1995) Organization of orosensory responses in the nucleus of the solitary tract of rat. J Neurophysiol 73:2144-2162.

Wu A, Dvoryanchikov G, Pereira E, Chaudhari N, Roper SD (2015) Breadth of tuning in taste afferent neurons varies with stimulus strength. Nat Commun 6:8171.

Xie K, Fox GE, Liu J, Lyu C, Lee JC, Kuang H, Jacobs S, Li M, Liu T, Song S, Tsien JZ (2016) Brain Computation Is Organized via Power-of-Two-Based Permutation Logic. Front Syst Neurosci 10:95.

Yamamoto T, Yuyama N, Kato T, Kawamura Y (1985a) Gustatory responses of cortical neurons in rats. III. Neural and behavioral measures compared. J Neurophysiol 53:1370-1386.

Yamamoto T, Yuyama N, Kato T, Kawamura Y (1985b) Gustatory responses of cortical neurons in rats. II. Information processing of taste quality. J Neurophysiol 53:1356-1369.

Yokota T, Eguchi K, Hiraba K (2011) Functional properties of putative pyramidal neurons and inhibitory interneurons in the rat gustatory cortex. Cereb Cortex 21:597-606.

